# A pipeline for the reconstruction and evaluation of context-specific human metabolic models at a large-scale

**DOI:** 10.1101/2021.07.22.453372

**Authors:** Vítor Vieira, Jorge Ferreira, Miguel Rocha

## Abstract

Constraint-based (CB) metabolic models provide a mathematical framework and scaffold for *in silico* cell metabolism analysis and manipulation. In the past decade, significant efforts have been done to model human metabolism, enabled by the increased availability of multi-omics datasets and curated genome-scale reconstructions, as well as the development of several algorithms for context-specific model (CSM) reconstruction. Although CSM reconstruction has revealed insights on the deregulated metabolism of several pathologies, the process of reconstructing representative models of human tissues still lacks benchmarks and appropriate integrated software frameworks, since many tools required for this process are still disperse across various software platforms, some of which are proprietary.

In this work, we address this challenge by assembling a scalable CSM reconstruction pipeline capable of integrating transcriptomics data in CB models. We combined omics preprocessing methods inspired by previous efforts with in-house implementations of existing CSM algorithms and new model refinement and validation routines, all implemented in the *Troppo* Python-based open-source framework. The pipeline was validated with multi-omics datasets from the Cancer Cell Line Encyclopedia (CCLE), also including reference fluxomics measurements for the MCF7 cell line.

We reconstructed over 6000 models based on the Human-GEM template model for 733 cell lines featured in the CCLE, using MCF7 models as reference to find the best parameter combinations. These reference models outperform earlier studies using the same template by comparing gene essentiality and fluxomics experiments. We also analysed the heterogeneity of breast cancer cell lines, identifying key changes in metabolism related to cancer aggressiveness. Despite the many challenges in CB modelling, we demonstrate using our pipeline that combining transcriptomics data in metabolic models can be used to investigate key metabolic shifts. Significant limitations were found on these models ability for reliable quantitative flux prediction, thus motivating further work in genome-wide phenotype prediction.

**Author summary:** Genome-scale models of human metabolism are promising tools capable of contextualising large omics datasets within a framework that enables analysis and manipulation of metabolic phenotypes. Despite various successes in applying these methods to provide mechanistic hypotheses for deregulated metabolism in disease, there is no standardized workflow to extract these models using existing methods and the tools required to do so are mostly implemented using proprietary software.

We have assembled a generic pipeline to extract and validate context-specific metabolic models using multi-omics datasets and implemented it using the troppo framework. We first validate our pipeline using MCF7 cell line models and assess their ability to predict lethal gene knockouts as well as flux activity using multi-omics data. We also demonstrate how this approach can be generalized for large-scale transcriptomics datasets and used to generate insights on the metabolic heterogeneity of cancer and relevant features for other data mining approaches. The pipeline is available as part of an open-source framework that is generic for a variety of applications.

## Introduction

Over the last decades, systems biology has enabled the comprehension of the different layers of biological processes, enabling the interpretation of the data generated by several emerging high-throughput technologies. The recent growth in both the diversity and quantity of data generated by these technologies led to the need of novel approaches for data processing, analysis and modelling to take full advantage of these data. Genome-scale tools for genomics, transcriptomics or proteomics allowed scientists to generate new approaches for experimental design and analysis, with important tools such as Genome-Scale Metabolic Models (GSMMs) granting the ability of simulating cellular processes and infer its phenotype, saving time and costs [1].

Metabolism is one of the main fields of study for systems biology which saw a significant evolution. It is crucial for the study of cellular function and its disturbance may lead to several diseases, ranging from diabetes or hypertension to cancer [2, 3]. Some of these problems can be diagnosed from metabolite screenings in human blood or urine [4] that are being exploited to help the discovery of treatments for the aforementioned diseases [5, 6].

Cells can be viewed as a multi-layered system, and the representation of reactions and metabolites is not enough to understand the whole metabolism. Several efforts have been made in the past years to provide for the integration of other layers, such as the genome or transcriptome, leading to a better understanding of other challenges, such as determining the fluxes through metabolic reactions. This led to the generation of GSMMs, a tool that has been heavily used in metabolic engineering [7, 8] and study of metabolic diseases [9–12].

Recon1 [13] was the first human GSMM, released in 2007. Since then, other metabolic models, such as the Edinburgh Human Metabolic Network (EHMN) and Human Metabolic Reaction (HMR) were developed, with constant revisions and integration occurring with many contributions [9, 14–18]. However, the lack of standardization of certain properties, such as gene or reaction nomenclature, led to propagation of errors throughout the years. The most recent open-source GSMM, Human-GEM, integrates the knowledge from previous models and tries to solve the main problems found, resorting to a joint effort of the scientific community [19]. This model comprises 13,417 reactions, 10,138 metabolites, 4164 being unique ones and 3625 genes.

GSMMs’ potential for phenotype simulation can be attained through constraint-based modelling (CBM) methods, which allow for fast calculations over large algebraic models, under the assumption of a steady state (concentration of internal metabolites assumed to be constant over time), since most kinetic parameters for metabolic reactions are not known. A stoichiometric matrix *S* depicts the main structure of the model, with columns defining reactions and rows metabolites, where *S_ij_* represents the coefficient of the *i*-th metabolite in reaction *j*. The assumption of the steady-state can be expressed as: *S* · *v* = 0, where *v* is the flux distribution vector. In addition to these constraints, every reaction in the model has an upper and a lower bound, restraining the maximum and minimum amount of flux passing through it.

These constraints define an admissible space of flux distributions, i.e. the rate at which every metabolite is either produced or consumed for each reaction [20].

These models can be used to support the definition of linear optimization problems, setting their constraints, and allowing different formulations to be achieved through appropriate objective functions over the flux distributions (*v*). A common approach is to define an equation (pseudo-reaction) representing the growth of a cell, which will be maximized [21], thus defining Flux Balance Analysis (FBA), a linear programming (LP) problem [22]. Despite its limitations (gene regulation, signaling processes or metabolic regulation are not taken into account), FBA has been successfully used to assess wild-type phenotypes, but also the impact of gene knockouts on metabolism [23–25].

Due to its simplistic and flexible approach, FBA motivated several extensions to address some of its limitations and allow further analyses. One of these variants is the Parsimonious enzyme usage FBA (pFBA), a method that relies on the assumption that flux distributions that demand the lowest overall flux through the network and are quicker to grow are selected, leading to improvements in the assumption made FBA [26].

GSMMs enable the integration of several types of information, including distinct omics data. A relevant application of integrating these data is the generation of tissue/cell specific metabolic models [27], starting from a general GSMM, for example, the Human-GEM. Indeed, these general puprpose species-level models contain information for all of the known metabolic reactions present in all types of human cells. While this can be useful to understand some generic processes of the human metabolism, it may run short on what it may provide in specific cell types or tissues. Context-specific models can be particularly useful for researching cancer metabolism, since they have been shown to be able to simulate rapid growth, mutations in metabolic genes and the Warburg effect (aerobic glycolysis) [28].

Several methods were developed to build draft cell/tissue-specific metabolic models throughout the past years, taking as inputs a template generic GSMM and different types of omics data. Although these methods use different approaches, their final objective is to obtain a draft model and/ or a flux distribution which tries to match the omics data provided [28]. In this work, we will be focusing on the methods which retrieve a sub-model from the generic one, since we want to build representative models of cell lines rather than simulate a very specific condition.

As reviewed by Robaina-Estevéz and colleagues, these methods can be classified into three families, GIMME-, iMAT- and MBA-like. The main objective in the GIMME family is to try to reach fluxes obtained from the model consistent with omics data, while maximizing a Required Metabolic Function (RMF), such as growth. For the iMAT family, the objective is the same without the need of a RMF. For the MBA family, methods try to achieve model consistency according to a predefined core of reactions, which may come from literature or from the omics data (e.g. the most expressed genes) [29]. It should be noted that the choice of the algorithm has an impact on the quality of the reconstructed model [30–33]. Here, we will be using the FastCORE (from the MBA family) [34] and tINIT (both GIMME and iMAT families) [35] to assess the importance of the choice of method to reconstruct a cell/tissue-specific model.

The connection of expression data (transcriptomics or proteomics) to the GSMMs is possible due to the gene-protein rules included in the models, which contain information relating the genes encoding the enzymes associated with each reaction (if they exist), in the form of Boolean expressions. However, problems that may affect the accuracy of the reconstructed model, such as experimental and inherent biological noise, several possible platforms to obtain expression data, bias on the process of detection and non curated relationship between gene expression and reaction fluxes, are still a main concern [15].

Since the steps to reconstruct tissue-specific metabolic models address a combination of the aforementioned issues, it is necessary to establish a pipeline to integrate omics data (which will be referred to as “preprocessing” from now on). In the most common pipelines, there are three main steps to be fulfilled [36]. The first takes into account how to deal with reactions with isozymes, complexes and/or promiscuous enzymes, i.e. do not have a one-to-one relationship between gene and reactions (gene mapping). The second is the definition of a limit where a gene is considered either active or not (thresholding). The final one is which is the order of gene mapping and thresholding used in data integration. This study aims to provide a pipeline enabling to evaluate their importance.

Another important factor to take into account when reconstructing or simulating a metabolic model is the medium composition. By default, most of the GSMMs do not have a predefined composition, allowing the user to define which one to use. However, for most cases, it is difficult to discriminate all metabolites and their concentration, possibly leading to false predictions when the model is simulated [37].

It has been proven that different media can lead to changes in the phenotypic behaviour of the cells, how they respond to stress and change their epigenome or transcriptome. Specially in humans, there are some added components to the medium, such as fetal bovine serum (FBS) whose composition is not clear and add another unknown factor to how the model should behave on its presence [38].

As impactful as the other raised issues, the choice of an algorithm to reconstruct a tissue-specific model is another source of variability in the final model, mainly due to the differences on the reconstruction algorithm principle. As a validation method for the reconstructed models, they can be tested with a set of required metabolic tasks, such as production of lipids and vitamins [36]. Since not all tissues require the same set of tasks, some manual curation may be needed.

The aforementioned issues can pose challenges when reconstructing a large dataset containing different cell types. An example is the NCI-60 panel, used in the work of Richelle et al, which tried to overcome these issues with the integration of metabolomics data to improve the quality of the reconstructed models [36]. Another approach used by Robinson et al was the integration of enzyme abundance and kinetic information using a different framework, the Genome-scale model with Enzymatic Constraints using Kinetic and Omics data (GECKO), allowing the reduction of space of possible solutions. Even though these approaches could lead to an improvement of the accuracy and consistency of the models generated through algorithms for context-specific reconstructions, in both cases a small number of cell types were used. Lack of data for certain cell lines was one of the main limitations [19].

When considering the issues described above, we understand that there are some limitations that are troublesome to overcome, such as the lack of a standardization of preprocessing methods, lack of certain types of omics data to fully characterize cells’ metabolism (such as fluxomics) or even limitations of the methods themselves, like the assumption of a steady-state or missing information for certain metabolic reactions.

With this in mind, we developed a generic pipeline for context-specific model reconstruction, allowing to test several ways of performing data preprocessing, as well as to choose the algorithm used for model extraction and validation. In a first set of experiments, we applied this pipeline to generate multiple reconstructions of the MCF-7 breast cancer cell line using recent transcriptomics data and knockout screenings from the Cancer Cell Line Encyclopedia (CCLE) [39–41], as well as fluxomics and proteomics data from a recent work by Katzir et al [42]. We complemented the validation of those reconstructions with analysis of gene essentiality and compared various sets of parameters with robust classification metrics.

Through this initial analysis, we were able to identify the best-performing set of parameters, which were applied to a larger case study. So, in a second set of experiments, the best configurations were used to reconstruct models for the whole set of the CCLE cell lines (over 700). We validated the resulting reconstructed models with CRISPR gene essentiality screens. With this work, we aim to establish a pipeline to find the optimal tuning of parameters for a given dataset, to help to reconstruct a more insightful tissue-specific metabolic model. An important advantage of this work is the use of open-source software in all steps of the pipeline, unlike previous studies mainly using proprietary software.

## Materials and methods

The context-specific metabolic reconstruction (CSMR) pipeline employed in this work contains four essential steps: input preprocessing (1), context-specific reconstruction (2), refinement (3) and validation (4).

An overview of the work is present on the Figure 1.

**Fig 1.**
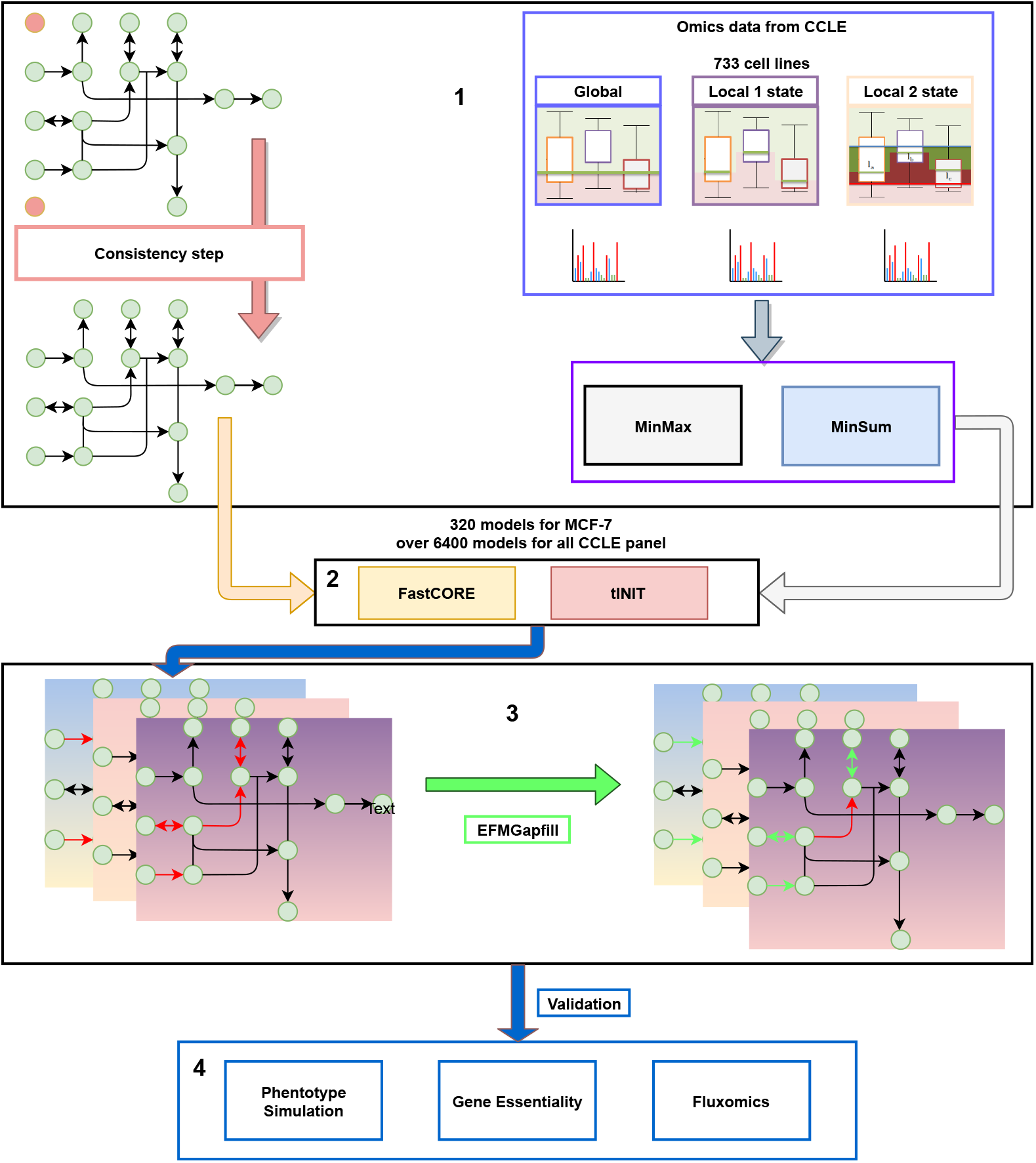
Overview of the context-specific metabolic reconstruction. In (1), we preprocess the input; first, we run the original model through a consistency step in order to remove some dead ends in the model; after, we preprocess the data from 733 cell lines through three different approaches to the threshold: global, local 1 state and local 2 state. In preparation of the reconstruction, we applied both MinMax and MinSum methods to include the gene information in the model and employed the FastCORE and tINIT algorithms (2). There were 320 reconstructed models for MCF-7 and over 6400 for the whole CCLE panel. Since this models may need refinement, all these models are subjected to a gapfill algorithm, EFMGapfill (3). Finally, the models are subjected several types of analysis, from phenotype simulation, gene essentiality to fluxomics (4).

### Input preprocessing

The inputs for any context-specific reconstruction always involve a template genome-scale metabolic model capable of yielding a non-zero flux distribution and only containing reactions capable of carrying non-zero flux, as well as a set of omics measurements integrated in the model via CSMR algorithms.

### Model preprocessing

We first ensure the model is consistent by identifying blocked reactions - whose maximum and minimum fluxes are null under open exchange conditions - with flux variability analysis, using the *find_blocked_reactions* function from the COBRApy package. We also remove any boundary metabolites - usually added to balance exchange reactions - prior to running any reconstruction, gapfill, or analysis method. This model must be feasible in steady-state conditions and must be capable of allowing flux through the biomass pseudo-reaction.

### Transcriptomics data as a proxy of enzyme activity

In this pipeline, we focus specifically on transcriptomics data that is mappable with the model’s gene associations, although some methods allow integration of other data types.

Similarly to previous approaches [19, 36, 43], we use transcriptomics data as a proxy for enzyme presence and flux activity from which we can calculate reaction activity scores (RAS) to serve as inputs for CSMR algorithms. These scores should ideally reflect whether a given reaction in the model is likely to be present in the context represented by the transcriptomics data. The work of Richelle et al. details the implications of several ways to infer RASs and provides several thresholding options [43], which we adapted as a part of our work. Out of the parameterization choices highlighted by the authors, we focused on varying the thresholding approach and GPR integration functions.

Using transcriptomics to characterize enzyme activity is not trivial, since the relationship between messenger RNA and protein expression is not fully understood, despite ongoing progress in quantifying both of these biological entities. However, Nusinow et al. have recently quantified the proteome for a subset of the CCLE panel and found a moderately positive correlation (mean Pearson c.c. = 0.48) between mRNA and protein abundance [44]. In this work, we assume a linear relationship so that RASs are calculated based on gene expression measurements.

### Scoring transcript activity from expression measurements

RNA-Seq technologies typically produce transcript-level measurements represented as proportions of the entire transcriptome. However, we intend, for the RASs used in this work, to obtain a positive or negative value relative to reference thresholds calculated across all samples. To this end, we process expression measurements into transcript activity scores (TASs) that can better represent this dichotomy.

We first define the concept of a global threshold, where the expression of all genes contribute. This is useful in filtering out transcripts whose expression is low or high enough for them to be assigned as inactive or active, respectively, with a high degree of confidence. However, this type of thresholding does not take into account the variability of measurements between transcripts. While originally available for microarray-based transcriptome quantification [45], a generic expression barcode for all cell types is hard to define and apply in RNA-Seq measurements, due to the different conditions in which experiments are performed.

A local thresholding approach can also be considered to mitigate the aforementioned problems. Rather than condensing the entire measurements into a single value, thresholding can be performed on a per gene basis, yielding a value for each gene independently. Similarly to the global thresholding approach, local thresholds can also be used as a reference for fold change calculations.

In this pipeline, both thresholds are calculated by first determining transcript-wise quantiles for various percentages (from 10% to 90%), yielding multiple sets of local thresholds, one for each percentage. To convert the latter to global thresholds, we use the mean value of a local threshold set to obtain a single representative value for the entire expression dataset.

After establishing appropriate thresholds, we can then combine them to establish rules that can be used to determine whether a transcript is active and calculate its TAS to better represent that activity level. We implemented an approach based on the work of Richelle et al [43], where transcript activity can be represented in two main states, namely:

- Inactive: transcript expression is inactive with a high degree of confidence - assigned when expression values are lower than a global lower threshold (*g_min_*). The expected TAS values will always be negative.
- Active: transcript expression is active with a high degree of confidence since its value exceeds a global upper threshold (*g_max_*). The expected TAS values are always positive.

This also implies the existence of an intermediate state of uncertainty for cases where transcript expression lies between the two thresholds. In these cases, we distinguish between active and inactive transcripts by comparing the transcript’s expression with its transcript-specific local threshold (*l*(*y*)) and the expected TAS value is constrained between −1 and 1, with its sign reflecting whether it is considered active.

We implemented three thresholding strategies based on the work of Richelle et al [43] with some minor changes to accommodate for the expected distribution of TAS values across multiple states. A global thresholding strategy implies a single global threshold to distinguish between active and inactive transcripts. An extension of this strategy, named local 1-state, includes local thresholding for transcripts that would otherwise be considered as inactive, and assigns TASs based on the ratio between expression and the transcript’s local threshold. Finally, we also included a local 2-state strategy defined by the usage of two thresholds and an intermediate state, as defined above.

TASs are then generated by calculating the ratio between measured expression values and an appropriate threshold which is chosen according to the state in which the transcript is assigned. A detailed description of the formulae used in each state for the three employed strategies can be found on Table 1.

**Table 1.** Functions used to convert transcript expression values into transcript activity scores, assuming *x* as a vector of expression levels for each transcript. The “Expression value” column represents the condition that values in *x* must meet for the corresponding reference threshold *t* used to calculate a ratio with the formula 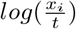. Finally, the range of TAS values for each condition is detailed on the last column. * In the local 2-state strategy, the TAS range does not start at 0 since 1 is added to the formula in the specific case when the expression value is greater than *g_max_*

| Strategy | Expression value | Reference threshold | TAS range |
| --- | --- | --- | --- |
| Global | $x \geq 0$ | $g_{max}$ | $] - \infty, \infty[$ |
| Local 1-state | $x \leq g_{max}$ | $g_{max}$ | $] - \infty, 0[$ |
| | $x \geq g_{max}$ | $l(y)$ | $] - \infty, \infty[$ |
| Local 2-state | $x \geq g_{max}$ | $g_{max}$ | $[1, \infty]^*$ |
| | $x \leq g_{min}$ | $g_{min}$ | $] - \infty, 0[$ |
| | $g_{min} \leq x < g_{max}$ | $l(y)$ | $[-1, 1]$ |

### Inferring reaction activity from transcript scores

The TASs from the aforementioned strategies are then converted to RASs using the gene-protein-reaction (GPR) rules provided with the model. GPR rules are Boolean expressions that describe, for a given reaction, which combinations of transcripts are involved with the synthesis of one or more enzymes capable of catalyzing it. These rules are often expressed or can be converted into disjunctive normal form, where multiple conjunctions (expressions with the AND operator) denoting the various enzymes or isoforms involved are bound by a disjunction (OR operator).

RASs must be presented as continuous scores and, thus, the Boolean operators in GPR rules must be replaced with numerical values. We replaced AND operators with a *minimum* function - an enzyme’s activity is limited by the lowest expressed transcript/subunit - while OR operations could be replaced with either *sum* or *maximum* functions. When using sum, we assume the reaction activity correlates with the combined activity of all enzymes and isoforms catalyzing it, while the *maximum* function equates reaction activity with the highest expressed enzyme’s score.

### Model reconstruction

#### Normalizing inputs for context-specific reconstruction algorithms

This step includes conversion of RASs into inputs accepted by the different CSMR algorithms, given their diverse nature, and we have implemented two alternatives to perform this conversion in our routines. Although our pipeline is generic, we have chosen the FASTCORE [34] and tINIT [46] algorithms for context-specific model reconstruction.

Methods such as tINIT, where scores mirror the reactions’ states as present or absent, can take RAS as input without any further processing. On the other hand, algorithms such as FASTCORE require a set of core reactions as input. In this case, a further threshold must be applied for these to be obtained. In our work, we emphasized the division between positive and negative scores to represent activity, and as such, core reactions are those with a RAS above 0.

#### Algorithm output and post-processing

With the inputs appropriately adapted, the output of each algorithm is always a binary vector *r* of size *n* (equal to the number of reactions in the template model), indicating reactions’ presence. Indeed, this vector includes Boolean flags indicating whether each reaction should be kept or removed in the context-specific model.

The models generated by the CSMR algorithms are then checked for consistency with expected phenotypes. For each of these models, we knockout (set lower and upper bounds to 0) reactions flagged for removal before to perform any simulation or analysis. We first check whether the model is capable of allowing non-zero flux through the biomass reaction, to ensure lethality can be tested. When growth medium formulations are available, we can additionally ensure that the model is feasible and capable of growth if the compounds present in growth media are the only ones allowed to be consumed. This is achieved by constraining exchange reactions that do not involve medium metabolites to only allow positive flux values, thus only allowing medium metabolites to be consumed by the model.

### Refinement

When the preliminary checks described above fail, gap fill approaches can be employed to infer sets of missing reactions that can expand the solution space and enable expected phenotypes.

#### Elementary flux mode-based gap filling approach

Gap filling was performed using a novel EFMGapfill approach, which was implemented in the *troppo* Python package. This algorithm leverages efficient elementary flux mode (EFM) enumeration algorithms to find minimal sets of active fluxes required for feasibility under a specific condition. Assuming a stoichiometric matrix *S* of *m* metabolites and n fluxes, the flux vector *v* and an identically sized vector *y*, and a set *K* of reactions available to fill gaps, the LP formulation employed in EFMGapfill can be defined as follows:

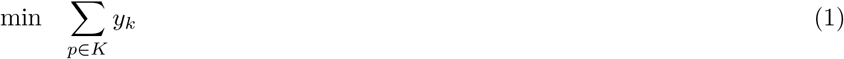

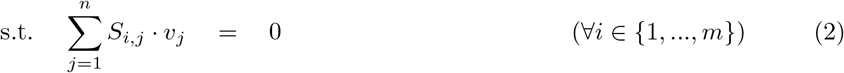

(LP1)

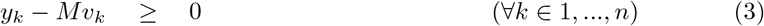

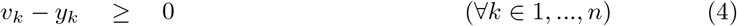

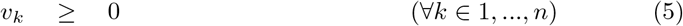

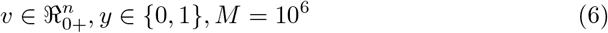

In the formulation represented in LP1, constraint 1 defines the steady-state constraint, similarly to other CB approaches, such as FBA. Constraints 2 and 3 associate the binary variables in *y* to the fluxes in the vector *v*. In this expanded solution space, the variables in *y* will hold a value of 1 if their associated flux in the vector *v* is greater than 1. Otherwise, both variables are set to 0. These variables and indicator constraints are then used to discretize fluxes into active and inactive states. The objective function is dependent on the set of reactions *K* available for the algorithm to add as a gap filling solution, although the objective is always to minimize the sum of a subset of the vector *y* whose indices are contained in *K*.

We adapt the input *K* according to a Boolean vector *r* (a set of reaction indices). *K* will have all reactions from the template model not included in *r*. The vector *r* is typically the output of a previously determined CSMR reconstruction. Furthermore, we also define and constrain an objective reaction *u* (usually the biomass pseudo-reaction) to always carry non-zero flux, representing a phenotype that is expected to be maintained upon tailoring the model to the subset of reactions in *r*.

We identified two possible gap filling scenarios that can be accomplished using this approach. In the first, we do not assume constraints on external metabolite exchanges and thus, we also exclude these reactions from *K*. The resulting solution from our gap filling approach is the smallest set of intracellular reactions not found in *r* that should be included so that the context-specific model is capable of carrying flux through *u*.

An alternative scenario may arise where the set of reactions *r* must not be manipulated, but the model still requires gap filling to predict growth. The growth medium, rather than the enzyme content of this model must be the target for manipulation. In this case, all intracellular reactions not in *r* must be constrained and exchange reactions must be split into forward and reverse reactions carrying flux in opposite directions. To find the minimal set of extracellular metabolites required for the model to carry flux through *u*, the set *K* must be defined as the set of reverse exchange reactions in the model.

### Validation

An important question arising from any CSMR process is ensuring the reconstructed models are capable of capturing the metabolic context of the cell or tissue, as represented by their corresponding omics measurements. Although literature review may reveal expected behaviours and phenotypes associated with the specific context to be modeled, a truly systematic validation of these models can only be achieved with large-scale datasets covering a wide range of measured biological entities. Such experiments should clearly point out the effect of perturbations that can be mapped onto the model on cell metabolism so that simulated fluxes become directly comparable. In this section, we describe how gene knockout screens and fluxomics can be integrated in our pipeline to validate these models.

#### Gene essentiality

Gene essentiality screens, such as those performed with CRISPR, provide a directly quantifiable measurement of the impact of gene deletions on cell viability, which can be modeled on CBMs through metabolic tasks [19]. The biomass objective function, included in most human models, groups most of these tasks’ demands by aggregating the necessary components for cell division and maintenance. Given the computational demand of checking multiple gene knockouts for each task and each omics sample, we will focus on predicting lethal gene knockouts using the biomass objective function as a measure of cell growth.

The CBM workflow used to predict essential genes uses GPRs to determine the set of reactions to exclude given a knocked-out gene *g*. To this end, we first obtain a mapping *ω*(*g*, *r*), which evaluates the GPR expression of reaction *r* with every gene marked as active, except for *g*. To apply the gene knockout, we must first determine the set *K* = {*r*|*r* ∈ *R*, ∀¬*ω*(*g*,*r*)}, which identifies the reactions that are disabled upon deletion of g; then, we set the lower and upper bounds of each reaction in *K* to 0. Adding these constraints to the model, the simulation can be run using FBA, yielding predicted growth rates for each gene deletion.

Finally, flux distributions resulting from gene knockouts can be evaluated. It is useful to always compare predicted mutant growth rates with wild-type levels. We considered several growth rate thresholds based on the wild-type value to represent viability, although previous studies have considered growth rates below 0.1% of the predicted wild-type rate to imply lethality. Additionally, infeasible solutions are considered as non-viable. We then discretize each gene knockout’s result as essential or non-essential and compare them with the experimental screening.

We used Matthews’ correlation coefficient (MCC) to assess the predictive ability of our models. The multiclass definition of MCC as implemented in the *scikit-learn* package is presented on Equation 7, assuming a generic classifier to predict

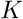

classes, *t* as a vector with the amount of true positives and *p* the vector with the amount of predictions each class *k*, while *c* is the total amount of true positive samples for all classes and *s* is the number of samples.

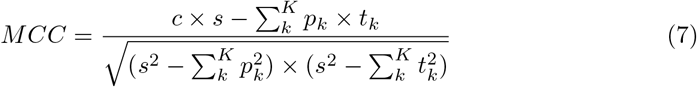

#### Predicted fluxes

Alternatively, flux distributions obtained from the model using an appropriate phenotype prediction method can be directly compared with experimentally measured fluxes, obtained from techniques such as isotope labeling coupled with metabolic flux analysis. In this work, we employ parsimonious enzyme usage flux balance analysis (pFBA) to predict phenotypes using our context-specific reconstructions. We have chosen this method since it requires no prior knowledge and reduces the admissible solution space of FBA by assuming cells not only attempt to achieve the predefined cell objective, but also minimise the overall sum of metabolic fluxes to do so.

A key limitation in using CB models to predict flux values is the lack of reliable measurements for substrate uptake fluxes. This directly influences the predicted growth rate and intracellular fluxes. A more reliable comparison can be made by discretizing flux values into three classes: *forward active,* if the flux is positive, *reverse active* if it is negative (flux is active, but carried in the reverse direction), or *null* when there is no flux. Although less precise, this discards the usage of experimentally measured external metabolite consumption rates. The model’s predictive ability can then be ascertained by using metrics suitable for multiclass predictive models such as Matthews’ correlation coefficient or weighted F1 scores.

### Flux analysis

Models reconstructed using our pipeline yielded flux distributions obtained from pFBA that were used for further analyses. Before applying decomposition methods, statistical tests or using these data for classification tasks, we first scaled flux values to avoid numerical issues. To achieve this, we transformed the entire dataset by applying a sigmoid function *s*(*x*) (Equation 8) that maintains flux signs, but brings very large values closer. Standardization was not performed as keeping the flux sign intact allows for proper interpretation of these values regarding alternative flux modes associated with the same reaction.

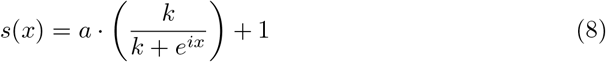

Relevant fluxes were selected before using supervised or unsupervised algorithms by eliminating fluxes with low variance. Furthermore, we also select an arbitrary number of features ranked by their significance in explaining the variance of the data relative to a discrete clinical feature using one-way ANOVA tests.

One of the methods used to analyse predicted fluxes is Principal Component Analysis (PCA), which we used to further reduce the high-dimensionality of the metabolic model’s solution space. We also inspected principal component loadings to identify groups of fluxes that were relevant with the clinical features in the biological samples from which the models were reconstructed.

Finally, we also used predicted fluxes to train supervised learning classifiers. We used Random Forest classifiers with varying number of Decision Tree estimators. K-fold cross-validation (CV) was used to assess the classifiers’ predictive performance using Matthews’ correlation coefficient as our metric. In some instances, we trained classifiers using several pFBA flux distributions for the same cell line. To avoid the inclusion of models from the same cell line in both training and testing folds, we have implemented a custom k-fold CV routine that splits datasets by cell lines rather than by individual flux distributions.

### Software availability

The software featured in this work was developed using the Python programming language. Although compatibility between language sub-versions should not cause any problems, we recommend using Python 3.6 and above. The entire source-code to perform all steps of the pipeline featured in this work is accessible through the GitHub repository at https://github.com/BioSystemsUM/human_ts_models/. The packages *cobamp*, *troppo*, *cobrapy*, *pandas*, *seaborn*, *scikit-learn* and *matplotlib* libraries are required to replicate the results and analysis featured in this work.

A significant part of our model reconstruction pipeline has been implemented using the *troppo* framework [47], developed in-house but freely available for the community. This software package provides an environment for omics data processing and subsequent integration with constraint-based metabolic models. This software is structured around two main parts: the omics layer handles data parsing, labeling and normalization, as well as mappings to previously loaded constraint-based metabolic models; the reconstruction layer contains routines to easily adapt omics inputs into appropriate reaction-level scores and run context-specific model reconstruction algorithms using novel implementations of existing methods.

We also used the *cobrapy* package [48] to read genome-scale metabolic models in the standardized Systems Biology Markup Language (SBML) format, manipulate their content and predict phenotypes using pFBA. The IBM^®^ ILOG^®^ CPLEX^®1^ (version 12.8) solver was used for all CB analysis and CSMR methods involving linear programming optimization problems, with or without mixed-integer constraints. Some parts of the omics data processing pipeline were performed using the *pandas* package. These routines have been generalized and included in the source-code of this work as auxiliary functions, although most parts of the input preprocessing pipeline are fully accessible through *troppo*.

The remaining parts of the context-specific model reconstruction have also been implemented in several components of *troppo*. Both fastCORE and tINIT algorithms used in this work were run using in-house implementations, which had been validated in a previous work [47]. The EFMGapfill approach is a novel addition to this software package and was implemented using an in-house implementation of the k-shortest EFM enumeration already available as part of *cobamp* [49]. This package was also used to run these routines with multiprocessing support whenever applicable.

The plots featured in this work were generated using the *matplotlib* and *seaborn* libraries.

## Results

Our first study evaluated the influence of different input processing methods on model reconstruction, using the MCF7 breast cancer cell line as a case study. We validated the predictive ability of each parameter setup through a comparison of predictions from the reconstructed models with expected phenotypes from gene deletion screens and fluxomics measurements. This cell line was chosen due to its common use in many previous studies, from which a large quantity of knowledge and omics data can be accessed.

In a second stage, using knowledge from these MCF7 models, we selected the best performing preprocessing options for each algorithm, and reconstructed various models for all cell lines, validating them with gene essentiality predictions. In the absence of fluxomics measurements, we also assessed whether such models could be used to generate relevant information for other tasks by using the result of several pFBA simulations as features for supervised machine learning approaches.

### Case-study setup

We used the Human-GEM (version 1.5.0) genome-scale metabolic reconstruction as our template model, stemming from a recent effort by Robinson et al. to provide a consensus metabolic model for Homo sapiens [19]. The model consists of 13417 reactions associated with a total of 3625 genes and 4164 unique metabolites, integrating knowledge from previous reconstructions. The model and auxiliary reaction and metabolite tables were downloaded from the corresponding version release on the GitHub repository at https://github.com/SysBioChalmers/Human-GEM.

The experimental data used in this work was obtained from two different sources. The Cancer Cell Line Encyclopedia provides RNA-seq transcriptomics data for over 56000 genes across 1270 unique cell lines. These expression values are represented in transcripts per million (TPM) and are already pre-processed using standardized GTEx pipelines. TAS calculations were performed across the entire dataset, although the only integrated scores were those whose associated genes were mapped to the template metabolic model.

These datasets are complemented with the Achilles dataset, characterizing lethal effects of over 18000 gene knockouts through CRISPR experiments [40, 41]. Gene essentiality scores from this experiment were generated using CERES [41] and processed further until these scores are normalized so that −1 and 0 represent, respectively, the median essential and non-essential gene knockout effects. For this work, we considered five essentiality thresholds evenly distributed across the range between −1.5 and −0.5. As of the first quarter of 2020, 739 cell lines had been included in the Achilles dataset [50], which we then selected as the candidates for our large-scale model reconstruction effort. Gene/transcript nomenclature was converted using the latest HUGO Gene Nomenclature Committee approved symbol mappings whenever needed [51].

Fluxomics measurements for MCF7 cell lines were obtained as part of the dataset used in the analysis of the work of Katzir et al. [42], where time-series LC-MS metabolomics were used to estimate the rates of 44 reactions in three growth media conditions. Despite being originally mapped to the reactions in the Recon 1 GSMM, we processed the data and matched these flux measurements and reaction directionality with the Human-GEM template model.

### Reconstruction of MCF7 cell line models

We first reconstructed models of the MCF7 cancer cell line by considering every possible combination of the parameters displayed on Table 2, excluding invalid combinations. In addition to these models, we included a MCF7 cell line reconstruction featured in the work of Robinson et al. as a baseline comparison [19].

**Table 2.**
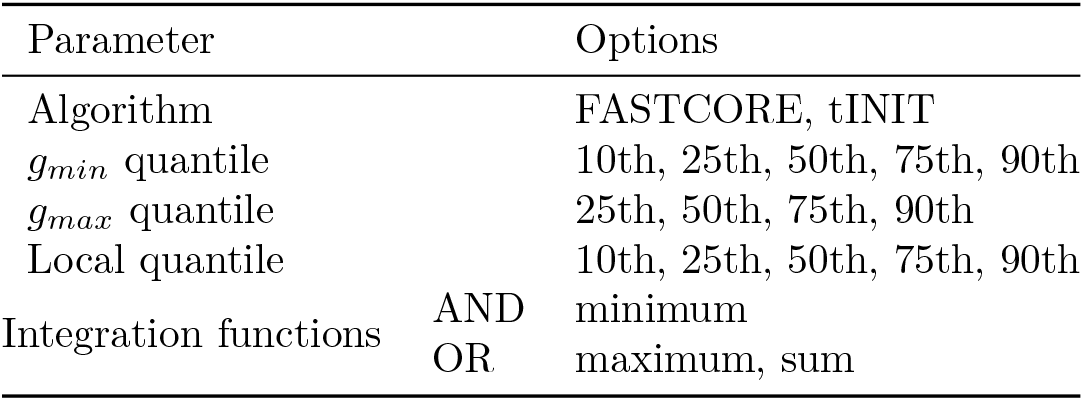
Required parameters for model reconstruction and possible options from which to choose from (separated by commas).

We obtained 320 models from this reconstruction effort and assessed their ability to correctly predict essential genes and flux activity. To understand the influence of parameterization on the models’ performance, we observed the distribution of values across multiple parameter options and evaluated parameter importance numerically using a linear regression.

#### Gene essentiality predictions

The results summarized on Figure 2 show that global thresholds have a greater impact on gene essentiality predictions, since the local 1-state strategy, which places a greater emphasis on local thresholding leads to worse gene essentiality predictions. Additionally, the average of all models reconstructed using the global thresholding strategy is close to that found in local 2-state models.

**Fig 2.**
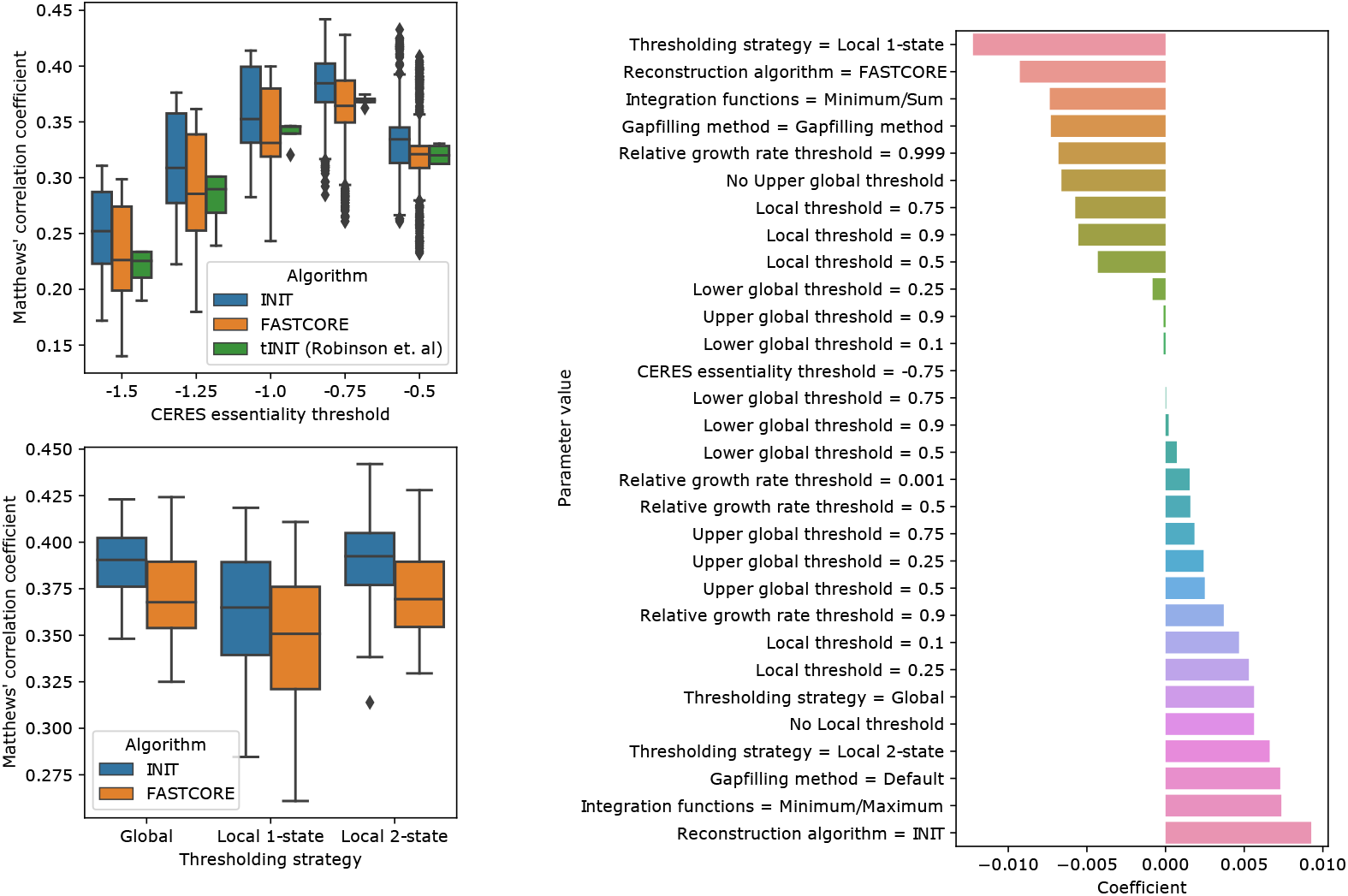
Overview of the influence of parameterization on the models’ performance when predicting essential genes as determined by their MCC. **Bottom left**: MCC value distribution for each thresholding strategy (horizontal axis) and algorithm combination (coloured box and whiskers). **Top left**: MCC value distribution for each CERES score threshold (horizontal axis) and algorithm combination (coloured box and whiskers). **Right**: Linear coefficients for each individual parameter value on a regression model aimed at predicting MCC values. Each parameter variable was one-hot encoded as multiple binary variables.

Despite this similarity when comparing the average of all models for each strategy, the best predictions were achieved using the local 2-state strategy, which is a clear indicator that a combination of both thresholding approaches are useful to estimate RASs. Although the performance achieved using the local 2-state strategy could be attributed to the usage of two (rather than one) global thresholds, we observed that the *g_min_* parameter has a negligible influence on the models’ predictive performance, which supports our claim that, in fact, both local and global thresholds have a positive effect when combined into the same TAS calculation strategy.

We were also able to infer some of the properties associated with the dataset, where *g_max_* and local thresholds at the 25th percentile seem to have the most positive effect on predictive ability in all models. Although the CERES score threshold representing the median essential gene knockout was set at −1, our models show slightly increased predictive power at −0.75.

Aside from data preprocessing related parameters, we have found the best parameter combinations are to use the tINIT algorithm in conjunction with the maximum function as replacement for the AND operator. We also observed that refining the model with EFMGapfill to allow growth using only the defined growth media metabolites as substrate did not result in better gene essentiality predictions.

#### Flux activity predictions

We also performed a similar assessment on the ability of our models to correctly predict reaction activity and directions for the MCF7 cell line under three growth medium compositions with associated fluxomics measurements. For each parameter combination, we generated three corresponding predictions using the growth medium as an additional flux constraint and calculated the MCC between the measured and predicted flux activities.

In Figure 3A, we can see that most parameters affect flux and essentiality predictions similarly. The best performing strategies are still those based on global and local thresholding with 2 states. In both algorithms, we also observed that constraining nutrient uptake to the metabolites that could be matched with the growth medium led to higher predictive power.

**Fig 3.**
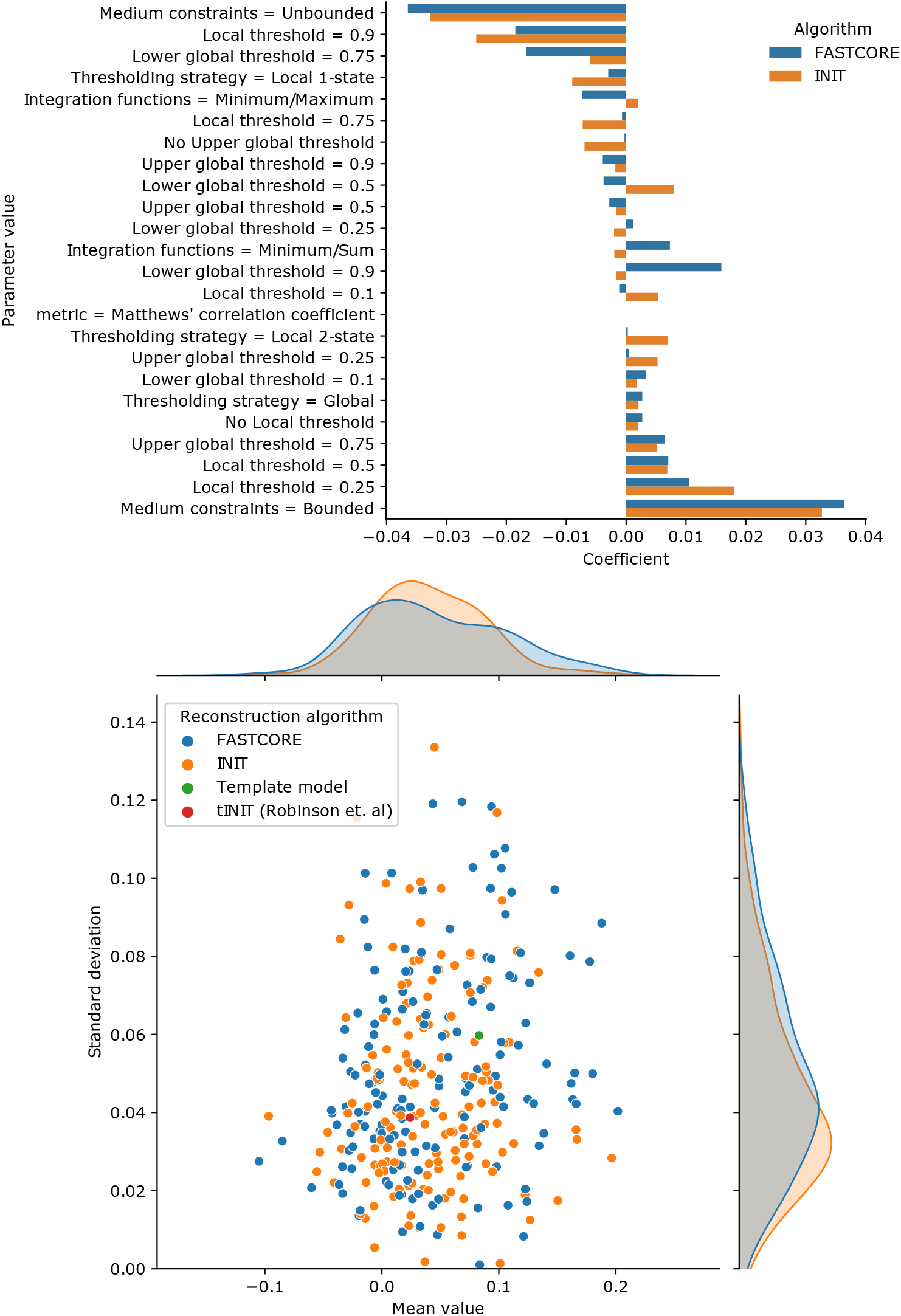
Overview of the influence of parameterization on the models’ performance when predicting flux activity and using MCC as the evaluation metric. **Top (A)**: Linear coefficients for each individual parameter value on a regression model aimed at predicting MCC values for each parameter combination. Each parameter variable was one-hot encoded as multiple binary variables. **Bottom (B)**: Relationship between average MCC value and standard deviation for each group of 3 simulations (conditions) that make up a single parameter combination. Different colors represent different algorithms and/or baseline comparison models.

The relationship between average MCC and its standard deviation is shown in Figure 3B where we can observe that FASTCORE reconstructed models were able to reach the highest correlation with the experimental fluxomics data, although they are more sensitive to parameterization. INIT, on the other hand, showed higher average MCC values across all parameter combinations with less dispersion and yielded models that rank closer when evaluated with this metric. We also compared our models with a baseline MCF7 cell line model featured in the work of Robinson et al. [19], which ranks significantly lower than our best FASTCORE and INIT models.

Overall, we did not significantly improve the predictions when considering a direct comparison of active fluxes between simulation and experimental quantification, although we identified some key reconstruction parameters that influence the context-specific model’s performance.

FASTCORE appears as the more consistent tool to extract context-appropriate sets of reactions from a template model. Due to its low computational demand, these reconstructions can be repeated with alternative parameters or to sample large amounts of models. tINIT, on the other hand, shows great potential to yield high-quality context-specific models, but thresholding parameters seem to heavily affect predictive ability.

### Large-scale metabolism reconstructions of cancer cell lines

We used 10 of the highest scoring parameter combinations from the MCF7 cell line case study to reconstruct the entire panel of cell lines available in CCLE with associated gene knockout effect screens (n=739). A similar reconstruction pipeline was employed in this larger case study, although we did not perform gap filling relative to the growth medium, due to heavy computational demand and an expected negative impact in phenotype predictions.

#### Predictive performance assessment

The results summarized on Figure 4 depict our large-scale results which show similar predictive accuracy to those reconstructed for MCF7 cell lines. There are slight differences in gene essentiality prediction performance between the 10 selected parameter combinations, with tINIT models reconstructed displaying slightly higher scores. In all of these scenarios, the selected pipeline parameterization choices improved gene essentiality predictions, when comparing with the models reconstructed in the work of Robinson et al, which asserts the importance of using more complex scoring strategies involving global and local thresholds.

**Fig 4.**
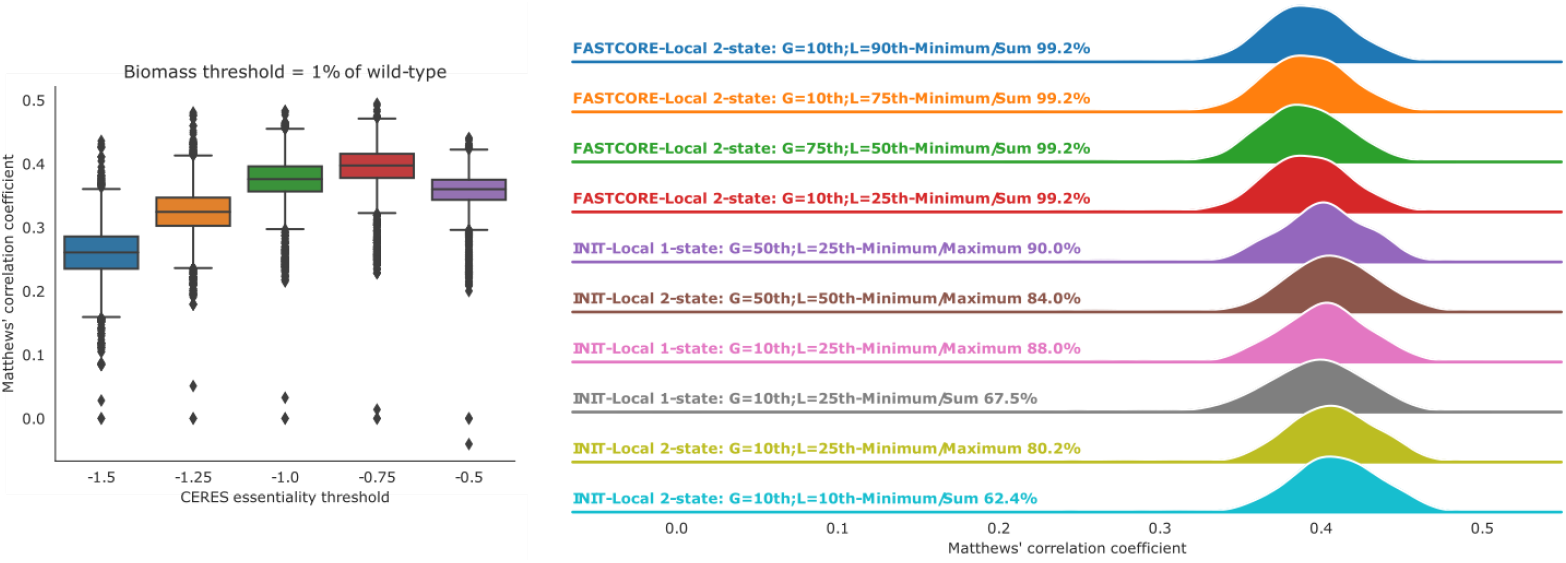
Overview of the predictive capability of the models reconstructed for each CCLE cell line. **Left**: MCC value distribution for all models in each lethal gene effect threshold. **Right**: Distribution of MCC values for each parameter combination selected for large scale reconstruction of CCLE models. The percentage value at the end of each thresholding strategy description represents the proportion of cell line models that could be successfully reconstructed.

The CERES score characterizing gene essentiality in the CRISPR experiments is undoubtedly the parameter that affects these predictions the most, with the best results obtained at a threshold of −0.75. This finding along with the overall low MCC values found with our approaches can be due to several factors. On the one hand, the biomass equation is a generalized assumption of the metabolites needed for cell growth, and thus, is not tailored for each specific cell line. The lack of more exact constraints on model uptake also results in gene knockouts that either do not affect flux through the biomass pseudo-reaction or completely inhibit it, which by itself elicits the usage of a threshold since a direct correlation between CERES scores and growth can not be found using constraint-based models.

### Exploring metabolic variability in breast cancer

We used the models and their respective predicted fluxes to explore the metabolic heterogeneity among various breast cancer cell lines. To do so, we retrieved molecular subtype annotations from the DepMap repository and used PCA to project these flux distributions using reduced features and obtain relevant information on the flux patterns present in different breast cancer subtypes. These results are summarized on Figure 5.

**Fig 5.**
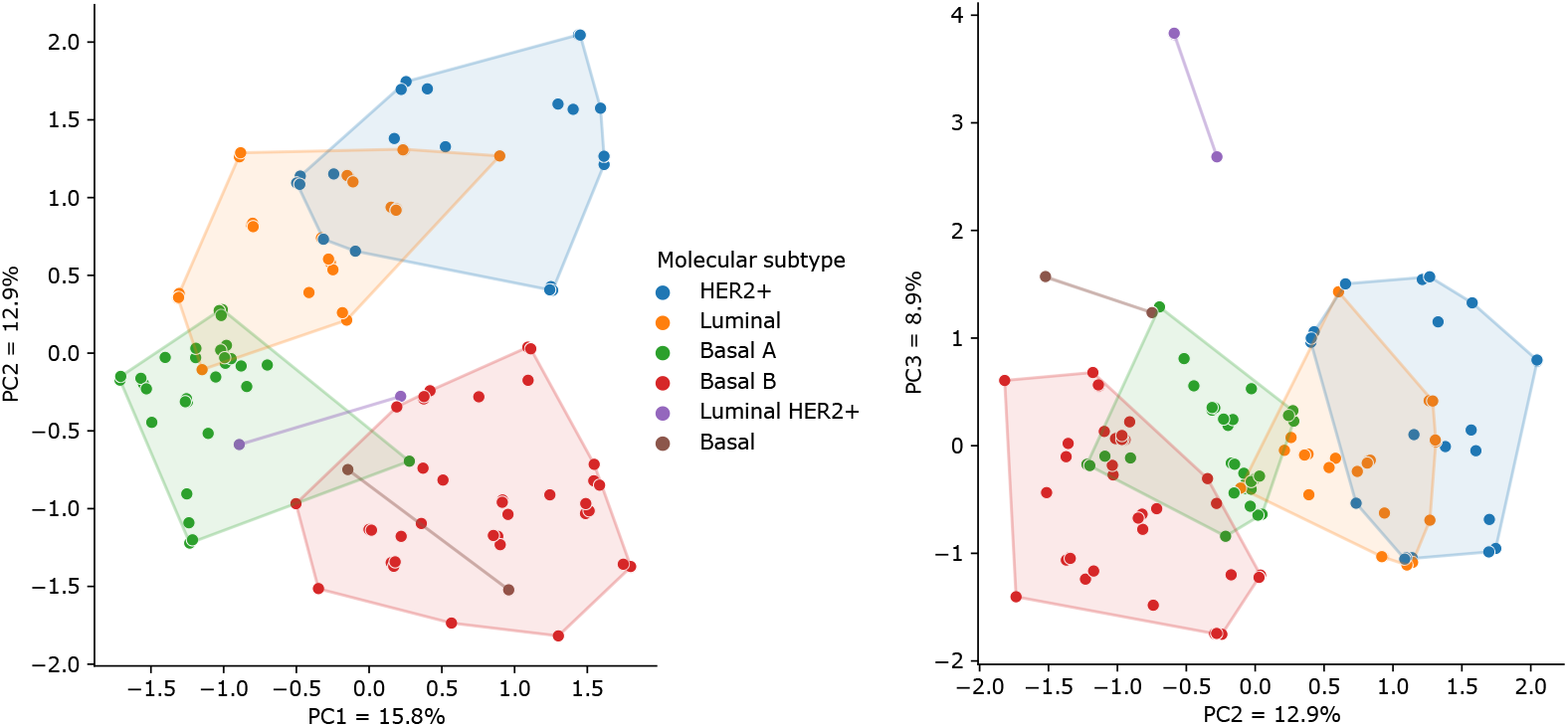
Cell line models reconstructed for breast cancer cell lines and projected in lower dimensions through the usage of PCA. The left figure shows the first PC against the second, while the right figure displays the second PC against the third, in the horizontal and vertical axes, respectively.

The subset of models belonging to breast cancer were extracted and the 200 fluxes with most statistically significant differences among subtypes were used to decompose the dataset. We included all reconstructed models, regardless of their parameters, and reduced the latent space to 3 principal components (PCs) capable of explaining 33% of the observed variance. The loadings of each principal component were obtained, with each flux being summarised in their corresponding pathways by calculating the average of the absolute ratios between the weights of each flux and the maximum observed values in the PC loadings.

Firstly, we can conclude that the decomposition of predicted fluxes can adequately distinguish between major molecular subtypes of breast cancer. The second PC (PC2) marks a good distinction between basal and luminal cell lines and correlates with reported prognosis and aggressiveness [52], with luminal BCs with better prognosis assigned to positive values as opposed to basal BCs which appear in this PC as negative values.

Lipid metabolism is typically deregulated in breast cancer and we were able to identify changes in fatty acid synthesis, with long-chain fatty-acid CoA ligase (ACSL1) and fatty acid desaturase (FADS) activity, as well as increased arachidonic acid production negatively correlated with the values in PC2. ACSL1 in particular has been reported as being transcriptionally upregulated in several breast cancer subtypes [53], and although it is not specific to basal BC, its expression has been shown to negatively impact survival rates. Moreover, previous studies have also shown that arachidonic acid promotes tumor cell migration in a basal B BC cell line [54], which further reinforces the association of PC2 with poor prognosis.

Another important finding was the negative correlation of PC2 with mitochondrial citrate carrier (SLC25A1) activity. This transporter plays a fundamental role in maintaining mitochondrial activity in high proliferating cells [55] and is highly expressed in triple-negative breast cancer (TNBC), corroborating the hypothesis that PC2 depicts a gradient of cancer aggressiveness.

### Predicted metabolic fluxes as relevant features

The lack of fluxomics data for the whole set of cell lines featured in DepMap does not allow to carry out a large-scale systematic comparison of the pFBA flux distribution predictions with experimental data. However, we set out to assess whether or not these predicted fluxes could be useful in predicting several clinical features associated with each sample. To do so, we established a supervised classification task, where the disease’s primary location would be predicted using various datasets, namely, (1) standardized expression values (TPM from RNASeq) using the entire gene set, as well as only those genes that can be integrated in the metabolic model, (2) TASs generated for each sample, (3) predicted fluxes (using pFBA over reconstructed models) and (4) the reaction presence (binary) from the CSMR algorithm outputs. Our results on this task are summarized on Figure 6.

**Fig 6.**
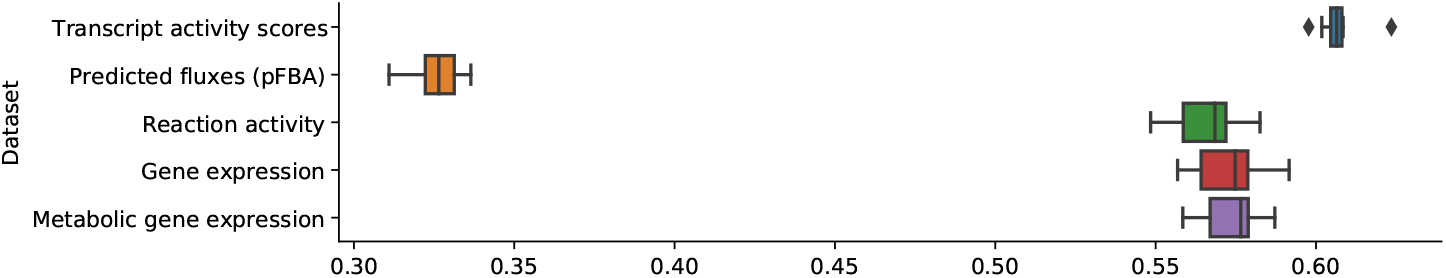
Distribution of average Matthews’ correlation coefficient values for cross-validated classifiers trained with various datasets

Classifiers trained with standardized transcriptomics data showed good relative performance (MCC mean=0.570, sd=0.038) with the subset corresponding to metabolic genes only slightly outperforming it. Processing these data and generating TASs slightly increased predictive capabilities (MCC mean=0.594, sd=0.030), which further justifies applying our preprocessing workflows before analysing and integrating omics data. However, the outputs of CSMR algorithms, namely, the presence or absence of each reaction resulted in models capable of predicting a cancer cell line’s primary site with an average MCC of 0.525 (sd=0.031). Furthermore, pFBA simulations resulted in even worse classifiers that could only reach an average MCC of 0.298 (sd=0.042).

Overall, our results show that context-specific model reconstruction and flux balance analysis approaches are not yet consistent enough for accurate quantitative flux predictions, as predicted metabolic fluxes by themselves did not appear to be relevant features for complex classification tasks.

## Discussion

We built upon several previous efforts to generate constraint-based models of differentiated human tissues, being capable of assembling a generic pipeline that can be useful in standardizing the process of integrating transcriptomics data into human metabolic models for the scientific community. Furthermore, this pipeline is available as part of an open-source software tool providing a generic framework for the implementation of context-specific model reconstruction tasks.

We were able to leverage large-scale multi-omics experiments with cancer cell lines and a state-of-the-art human metabolic reconstruction to generate meaningful models capable of capturing the metabolic diversity among, and within, multiple types of cancer. We were also able to validate the models using experimentally determined essential genes and fluxomics data.

The usage of decomposition methods to understand flux predictions allowed us to establish a link between metabolic phenotypes and breast cancer prognosis, and by making use of the interpretability of constraint-based models, we were also able to pinpoint key enzymes and metabolites associated with dysregulated growth. This elicits the potential for similar approaches to assist in contextualizing transcriptomics profiles into metabolic phenotypes, with the purpose of understanding the intricate mechanisms responsible for human diseases, especially for personalized medicine applications.

The availability of metabolomics and proteomics data still pales in comparison with RNA-Seq technologies used for transcriptomics quantification. As such, we have developed this work to only consider the latter omics type, and we argue that the reconstruction of models based on transcriptomics data results in computational tools that can be more easily adapted to a clinical setting since they do not rely on generating multiple omics datasets. However, we have also built the computational tools, namely *troppo*, in a way that these datasets can be easily integrated and used with appropriate methods.

Although encouraging, our results show the difficulty in closing the gap between experimentally measured and predicted fluxes. We argue that there is value in building representative models using gene expression alone, since the techniques used to obtain these measurements are far more ubiquitous and less costly. However, naturally, this lack of information implies some limitations when interpreting the model. This was evident when using model simulations to predict a cell line’s disease, where classifiers trained with these predictions displayed poor predictive performance.

In the absence of precise exo-metabolome uptake or secretion rates, CBMs in their original definition, are merely capable of predicting metabolic pathways on a discrete level, and thus, flux distributions must always be interpreted relative to a given original state or model context rather than assuming these fluxes are numerically comparable. A related challenge also appears when considering the biomass objective function, which is usually too generic to describe different tissue types, and hinders the ability for these approaches to generate meaningful models for cells whose metabolic objective is difficult to define.

Recent works that have incorporated exo-metabolite measurements [36], metabolic task protection and alternative formalisms to include more complex parameters [19], have reached better Pearson correlation coefficients with fluxomics measurements, although with smaller case studies. Another important aspect would be to expand the scope of constraint-based models to also include regulation and signal transduction enabling predictions of metabolic fluxes that can be contextualized with their corresponding regulators.

We must, additionally, acknowledge the importance of using a fluxomics data source for a reference cell line to serve as a basis for subsequent reconstructions, since it allows us to find ideal sets of parameters for larger-scale efforts. In this work, this led to a significant decrease in computational resource usage, as well as a better choice of parameters without exhaustive reconstructions.

The implementation of this complex pipeline in a modular framework allows for the usage of different methods that might fit a particular purpose. Previous works have reported the heterogeneity in outputs from various CSMR algorithms and our case study clearly shows that this choice impacts the type of phenotypes to predict and, as such, we extended *troppo* in such a way that reconstructing a context-specific metabolic model is a simple task, even for users with limited programming skills

## Acknowledgments

The authors thank the PhD scholarships co-funded by national funds and the European Social Fund through the Portuguese Foundation for Science and Technology (FCT), with references: SFRH/BD/118657/2016 (V.V.), SFRH/BD/133248/2017 (J.F.). This study was also supported by the FCT under the scope of the strategic funding of UIDB/04469/2020 unit.

## Footnotes

1 https://www.ibm.com/analytics/cplex-optimizer

